# Evolutionary dynamics under demographic fluctuations: beyond the effective population size

**DOI:** 10.1101/2025.04.12.648494

**Authors:** Kavita Jain

## Abstract

It has long been appreciated that changing population size can strongly affect the evolutionary dynamics and genetic diversity, and traditionally, the effects of demographic fluctuations are captured via an effective population size. However, as has been pointed out in previous studies, an effective population size does not always exist and can be defined only when the relevant time scales are well separated. A mechanistic understanding of the non-existence of effective population size is, however, missing, and a very few analytical results have been obtained when the relevant time scales overlap with each other. To address these issues, we consider a neutral, panmictic population whose size fluctuates in time, and for a general demography, we show that the correlations between the population size at different times preclude the existence of an effective population size. We then consider a specific demography in which the population size switches between two fixed values in a stochastic fashion, and formulate a diffusion theory that allows us to obtain exact results for fixation probability, fixation time and mean sojourn time of a mutant for arbitrary switching rates. We find that when the switching time and the time scale for the genetic drift are of the same order, these quantities differ substantially from the corresponding results predicted by a neutral model with an effective population size given by the harmonic mean of the population size, and discuss the implications of these results for neutral genetic diversity in changing environments.

## 1 Introduction

Natural environments are not static, and environmental changes over short term such as due to seasonal cycles [49] or in long term as, for example, arising from global warming [11] can result in changes in the population size and population fitness. A population must adapt to the changing environment to survive, and the genetic variation in an evolving population can be strongly affected by the changes in the population size [4] and/or selection [21]. Assuming that the environmental changes are deterministic, using branching process and diffusion theory, the impact of population bottleneck on neutral genetic diversity [35, 31], population growth on neutral fixation time [48] and neutral site frequency spectrum [50, 14, 25], and more general demographies on the fixation probability of a strongly beneficial mutation [9, 39, 44] have been investigated. Recent work have focused on understanding the effect of temporally varying selection on fixation probability [37, 43, 47, 6] and fixation time [1, 43, 24, 23] in populations of constant size, and how variation in both selection coefficient and population size affects various measures of genetic diversity [20, 2]. The effect of stochastic variation in selection coefficient [42, 13, 15] and fluctuating population size [30, 7, 53, 5] on the fixation process and site frequency spectrum have also been analyzed.

In general, evolutionary dynamics in changing environments involve several relevant time scales which can be broadly classified as those over which evolutionary forces (selection, genetic drift, mutation and migration) act when respective parameters (selection coefficient, population size, mutation and migration rate) are constant in time, and those that characterize the variation in them such as the time period of a cyclically changing parameter, decay time of correlations in a fluctuating parameter, etc.. In view of this complexity, it is useful to first consider the problem assuming that the relevant time scales are well separated; however, in general, such analyses do not provide an insight into the behavior at intermediate time scales. For example, for periodically varying selection coefficient, the fixation probability of a rare beneficial mutant in a panmictic, haploid population is twice the *initial* selection coefficient when selection changes very slowly and twice the *mean* selection coefficient for rapid changes, but it exhibits a minimum or maximum when the mean selection coefficient and the environmental frequency match (“resonate”) [6], while the corresponding fixation time changes monotonically between the above mentioned extreme limits [24].

The population dynamics at such intermediate time scales also raise important conceptual issues in a fluctuating environment. In a neutral, panmictic population, as considered here, the effect of stochastically varying population size has been studied within a coalescent framework; it is argued that an effective population size does not always exist and only when the time scales over which coalescent events occur and population size fluctuates are well separated, a (coalescent) effective population size can be defined [36, 22, 19, 41, 45]. However, the reason(s) underlying the non-existence of an effective population size have not been described. Furthermore, when an effective population size can not be defined, the patterns of neutral genetic diversity have been investigated mainly numerically [41, 34] (an exception is [8] in which branch length of neutral genealogies for arbitrary demographies are studied analytically).

Here, we consider a neutral model with fluctuating population size in forward time for which several quantities of interest can be solved exactly. The population is assumed to switch randomly between two sizes, that is, it instantaneously expands and contracts at exponentially-distributed times. This demography model has been considered in several previous work [16, 18, 40, 41, 17], but these studies were concerned with the question of the existence of an effective population size in fluctuating demographies. Here, we obtain exact expressions for the time-dependent fixation probability, conditional fixation time and sojourn time for a rare mutant, and study in detail how these quantities are affected by the rate of demographic fluctuations. Before proceeding to the specific model, we discuss how temporal correlations in general demographies preclude the existence of an effective population size.

## 2 Methods

### 2.1 General framework

In diffusion theory, the distribution of mutant allele frequency *p* in a neutral, panmictic, haploid population with changing population size *N*(*t*) obeys the following backward equation [26]:

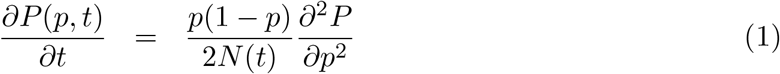

For appropriate boundary conditions, the solution of the above equation can be expressed as [26, 32]

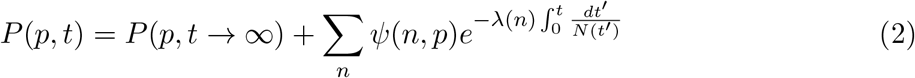

where the eigenfunction *ψ*(*n, p*) and eigenvalue *λ*(*n*) > 0 (of the neutral operator) are independent of the population size. For a deterministically changing population size or for a given trajectory of the fluctuating population size, expression (2) is sufficient to describe the desired distribution.

But for stochastically varying population size, one is often interested in the frequency distribution averaged over population trajectories. This requires one to find 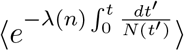 where ⟨…⟩ denotes the average with respect to the distribution of the entire population trajectory until time *t*, which is a formidable task. However, following (3.3b) of [29], one can express the mean of the exponential of the integral of a random variable (cumulant generating function) in terms of the exponential of the sum over time-integrated cumulants ⟨…⟩_*c*_,

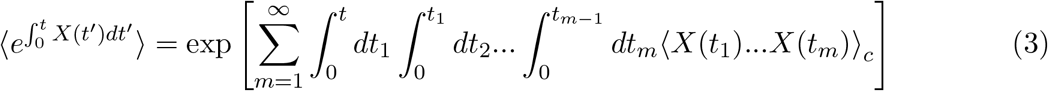

where *X*(*t*) is a random variable. Using this result, we can write

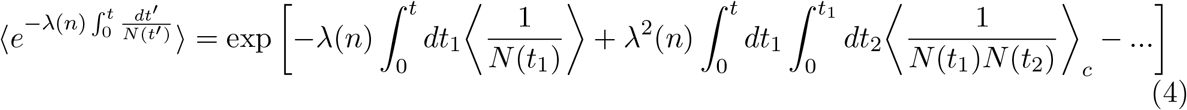

For constant population size *N*, as all the cumulants in the inverse population size vanish, the above expression reduces to 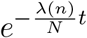 ; it is important to note that the exponent 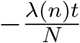 is *linear* in eigenvalue and time.

If one assumes that the population size are uncorrelated in time (that is, at each instant, they are unaffected by each other), all the correlation terms on the RHS of (4) vanish, and it reduces to 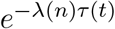 where 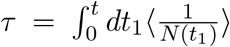 is, in general, nonlinear in time, but the exponent is still linear in the eigenvalue. If the (uncorrelated) population sizes are chosen from a stationary distribution, we can write

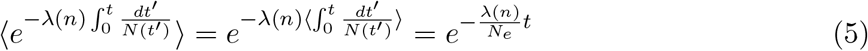

where the effective population size, *N*_*e*_ is the harmonic mean of the population size with respect to the stationary distribution. Thus, on comparison with the model with constant population size, we find that it is possible to treat the fluctuating population size problem as the one with a constant size *N*_*e*_ when the correlations are absent; mathematically, this correspondence exists because the exponential in (5) is linear in *λ*(*n*).

In biologically realistic scenarios, however, the population sizes are correlated in time and therefore, the exponent on the RHS of (4) has, in general, *nonlinear* terms in the eigenvalue *λ*(*n*). Two special cases [36, 19, 41] where the exponential in (4) is linear in the eigenvalue are: (i) if the population size fluctuates on time scales that are much smaller than the time scales on which random genetic drift acts, and (ii) if the population size changes slowly enough that it remains close to its initial value *N* (0) - in either case, we obtain (5), with *N*_*e*_ equal to the harmonic mean and *N* (0), respectively. Away from these extreme limits, the temporal correlations in the population size can not be ignored and one needs to go beyond the description in terms of an effective population size, and, in principle, all the cumulants in (3) need to be taken into account; in other words, an exact expression for 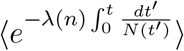 is required.

### 2.2 Model

We now consider a neutral, panmictic, haploid population with one biallelic locus whose total size fluctuates between two values, *N*_1_ and *N*_2_ > *N*_1_. In discrete time, the population size switches between the two states with a given probability, and at long times, this process reaches a stationary state (that is, the probability that the population size is *N*_1_ or *N*_2_ becomes time-independent). Then, at time *t* = 0, a single mutant replaces an individual in the wildtype population at its current size, and thereon the wildtype and mutant frequencies evolve via a neutral Wright-Fisher process defined as follows. At generation *t*, if the total population size is *N*_*i*_, *i* = 1, 2 and the number of mutant individuals is *n*, the mutant number in the next generation is obtained by binomially sampling 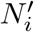 individuals with probability *n*/*N*_*i*_, where 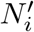 is the population size at generation *t* + 1. This process is continued until the mutant gets fixed at the current population size or becomes extinct; for further model and simulation details, refer to Appendix A.

### 2.3 Diffusion theory

The dynamics of the above model can be described analytically using a diffusion theory in which the state space has both continuous and discrete variables (see, for e.g., Section 3.4 of [12] for a detailed discussion). We first consider the dynamics of the population size: in continuous time, the population size switches from *N*_1_ to *N*_2_ at rate *γ*_1_ and *N*_2_ to *N*_1_ at rate *γ*_2_ - this is known as the random telegraph process (see, for e.g., Section 3.8.5 of [12]). In the stationary state of this process, the probability that the population size is *N*_*i*_ is given by 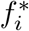,where

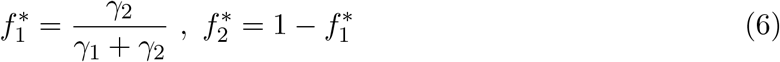

This result and other properties of the random telegraph process pertinent to the discussion are described in Appendix B.

We now consider the mutant population dynamics, and focus on the probability *P*_*i*_(*p, t*; *x*), *i* = 1, 2 that starting with mutant frequency *p* in population of size *N*_*i*_, the mutant population frequency is either 0 < *x* < 1 at time *t* or has reached the absorbing state frequencies, *x* = 0 or 1, by time *t*. These distributions obey the following coupled backward Fokker-Planck equations,

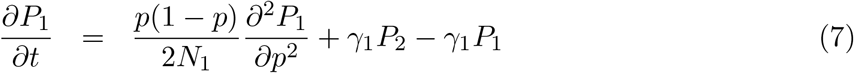

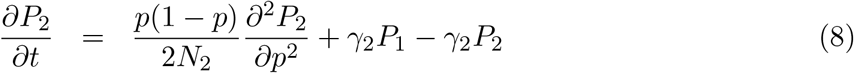

This is because in a small time interval *dt*, a change in these distributions occur either due to the usual neutral sampling in the population of constant size [first term on the RHS of (7) and (8)], or when the population size switches with the mutant allele frequency being *p*. The latter event means that the mutant initially present in the population of size *N*_1_ has frequency *x* at time *t* + *dt* either because the population size did not switch with probability 1 − *γ*_1_*dt* in the initial time interval *dt* and the mutant achieved the desired frequency in the time interval (*dt, t* + *dt*) with probability *P*_1_(*p, t*), or the population size switched from *N*_1_ to *N*_2_ with probability *γ*_1_*dt* in time *dt* and the mutant proceeded to frequency *x*, starting with frequency *p* in population of size *N*_2_. Thus the probability *P*_1_(*p, t* + *dt*) = (1 − *γ*_1_*dt*)*P*_1_(*p, t*) + *γ*_1_*dtP*_2_(*p, t*) which leads to the last two terms on the RHS of (7); on interchanging the subscripts, 1 and 2, we obtain the corresponding terms for *P*_2_(*p, t*) in (8).

## 3 Results

### 3.1 Fixation probability

If the mutant with initial frequency 0 < *p* < 1 eventually fixes, as the genealogies of all the individuals in the fixed population can be traced back in time to one of the initial mutants with equal probability, it immediately follows that the eventual fixation probability in a neutral population is *p*, irrespective of how the population size changes during the fixation process. But, as discussed below, the dynamics of fixation probability in a neutral population are affected by demography.

As we are interested in the probability that a single mutant arising in the population of size *N*_*i*_ fixes in population of either size by time *t*, equations (7) and (8) are subject to boundary conditions,

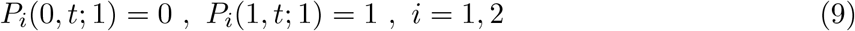

and the initial condition, *P*_*i*_(*p*, 0; 1) = 0 since the mutant population present in frequency *p* < 1 can not fix instantaneously. On averaging over the initial population size, we can write

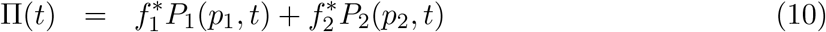

where, 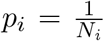,and 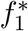 and 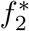 are given by (6). To check our diffusion theory, the results obtained by numerically solving (7) and (8) along with the above boundary and initial conditions were compared with those from discrete time simulations for various parameters, and found to be in good agreement (see Fig. S1).

In Appendix C, we show that equations (7) and (8) can be solved exactly, and find that

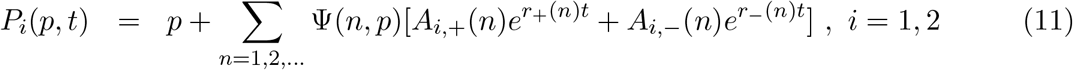

where, 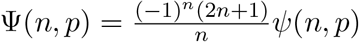,and *ψ*(*n, p*), *r*(*n*) and *A*(*n*) are given, respectively, by (C.4), (C.12) and (C.14). As *r*_±_(*n*) < 0, at large times, we obtain *P*_*i*_(*p*_*i*_, *t* → ∞) = *p*_*i*_, as argued at the beginning of this section.

To obtain an insight into the result in (11), we first note that while the genetic drift acts on a time scale that increases linearly with the population size [26], the population of size *N*_*i*_ remains in its state for time of order 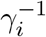 before switching to the other state. Thus the population size changes slowly compared to drift when 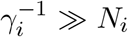 and rapidly for 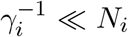. Using (D.4) and (D.9), we find that in these extreme cases, the fixation probability averaged over the initial population size can be approximated as

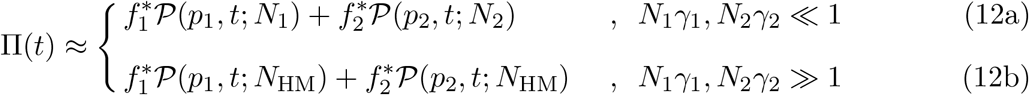

where,

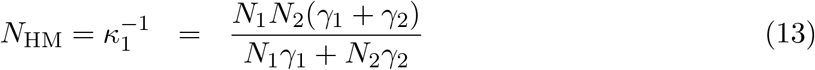

is the harmonic mean of the population size in the stationary state of the telegraph process (see Appendix B) and *N*_1_ < *N*_HM_ < *N*_2_. In equation (12), 𝒫 (*p, t*; *N*) denotes the fixation probability of the mutant by time *t*, when its initial frequency is *p*, in a population of constant size *N*, and given by 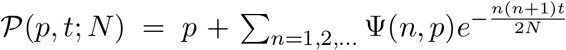 [26, 32]. Equation (12) states that the average fixation probability is the weighted mean of the fixation probability in a constant population whose size is the one in which the mutant arose if the environment changes slowly or given by the harmonic mean of the two population sizes (in the stationary state) in a rapidly changing environment.

Figures 1a and 1b show the average fixation probability Π(*t*) for various *N*_1_*γ*_1_ (keeping *N*_2_*γ*_2_ = *N*_1_*γ*_1_), and we find that Π(*t*) for small and large *N*_1_*γ*_1_ is indeed well approximated by (12a) and (12b), respectively. As discussed in Sec. 2.1, a description of the fluctuating size problem in terms of a constant population size requires the temporal correlations between population size to be negligible. If the mutant arises in the population of size *N*_1_ (say), the correlation between the inverse population size at two times, *t*_2_ and *t*_1_ > *t*_2_ are given by (E.6). For slowly fluctuating population size, this correlation function is proportional to 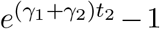 which is negligible for *N*_1_*γ*_1_ ≪ 1, if time is measured in units of population size; similarly, for rapidly fluctuating environment where this correlation function decays as 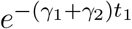,the population size are weakly correlated when the product of population size and switching rate is large. Away from these limiting cases, the population size are correlated in time, and as Figs. 1a and 1b depict, the fixation probability differs substantially from either extreme situation when *N*_1_*γ*_1_ ∼ 1.

**Figure 1:**
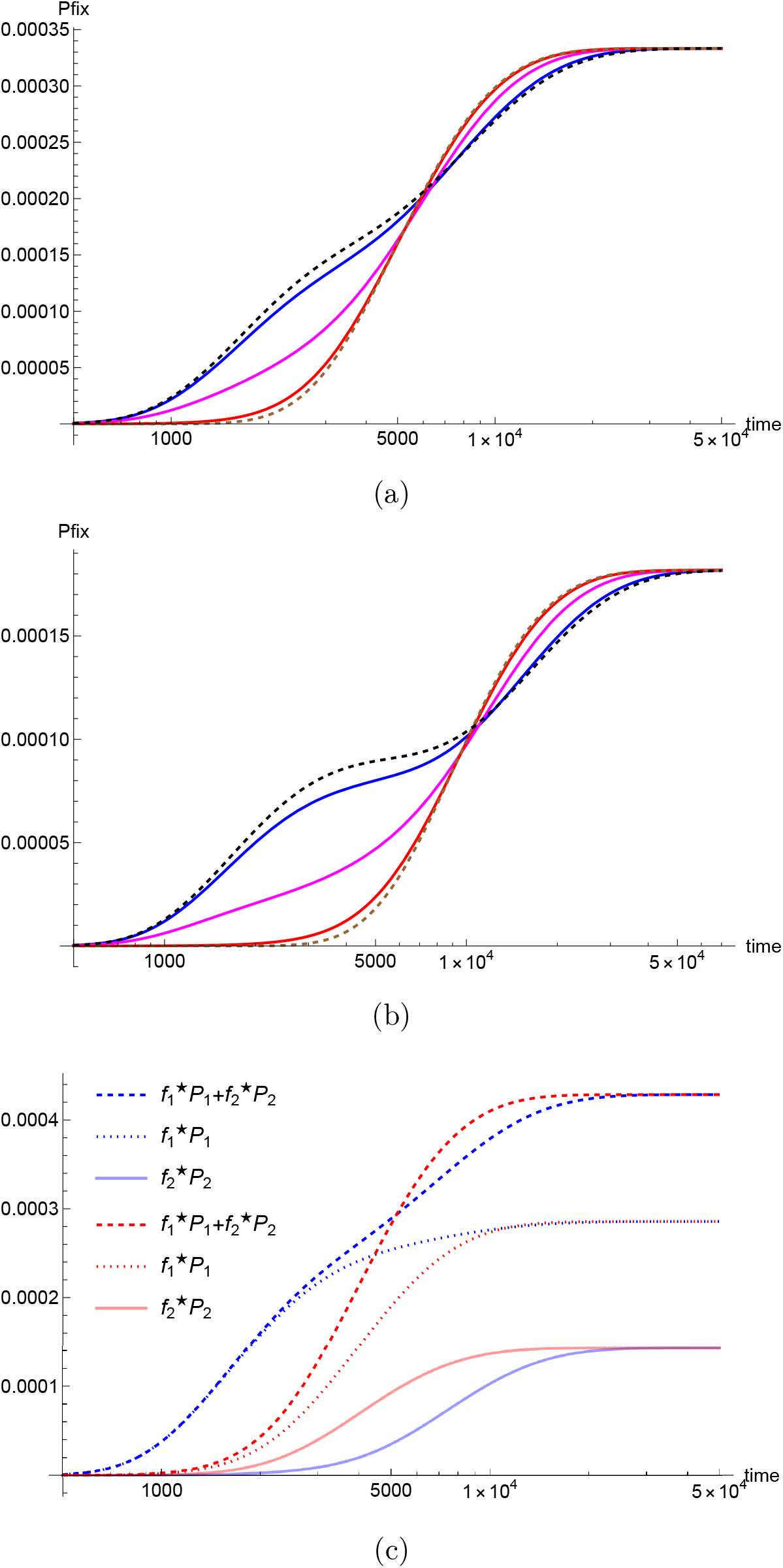
Average fixation probability, Π(*t*) of a single mutant for *N*_1_ = 10^3^, and (a) *N*_2_ = 5*N*_1_ and (b) *N*_2_ = 10*N*_1_ for *N*_1_*γ*_1_ = *N*_2_*γ*_2_ = 0.1 (blue), 1 (magenta), 10 (red), obtained by numerically solving (7) and (8). The black and brown dashed lines, respectively, show the expressions in (12a) and (12b). (c) Fixation probabilities as a function of time for *N*_1_*γ*_1_ = 0.1 (blue curves) and 10 (red curves) with *N*_1_ = 10^3^, *N*_2_ = 5*N*_1_, and *N*_2_*γ*_2_ = 2*N*_1_*γ*_1_. These curves are also obtained by numerically solving (7) and (8).

Figures 1a and 1b also show that at short times where the mutant population size is expected to be small and susceptible to loss due to genetic drift, the mutant is much more likely to fix if the population size changes slowly whereas at longer times, the fixation probability is mildly larger in a rapidly fluctuating environment. To understand these observations, we show the fixation probabilities Π(*t*) and 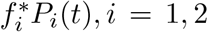 for slowly and rapidly fluctuating population sizes in Fig. 1c. As the time scale for genetic drift is proportional to the population size, the fixation probability in a population of size *N* is negligible for *t* ≪ *N* and reaches the eventual probability when *t* ∼ *N* . Then, at short times, in slowly fluctuating environment, due to the approximation (12a) and *N*_2_ > *N*_1_, the fixation probability 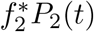 is negligible and the average fixation probability Π(*t*) is dominated by 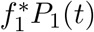,as shown in Fig. 1c. On the other hand, in rapidly fluctuating environment, due to (12b), the fixation probabilities 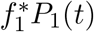 and 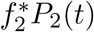 become non-negligible only at time of order *N*_HM_ > *N*_1_. As a consequence, at short times (*t* < *N*_1_), the average fixation probability Π(*t*) for slowly fluctuating size is larger than the corresponding result for the rapidly changing size.

At longer times (*t* ≫ *N*_1_), however, we need to compare the dynamics in populations of size *N*_2_ and *N*_HM_, and on repeating the argument above, we find the reversal in the qualitative trends. The long time behavior can be quantified by noting that the late time dynamics are governed by the smallest rate *r*_+_(1) in (11), which is given by

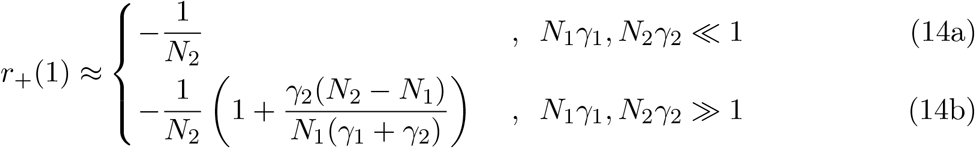

and shows that the long time behavior sets in at the largest time scale, namely, *N*_2_ in either case; this conclusion is indeed supported by Figs. 1a and 1b. Thus, the above equations show that in a rapidly changing environment, the fixation probability saturates sooner than in a slowly changing one if the two population size are substantially different, otherwise the long time behavior is essentially independent of *N*_1_*γ*_1_ (see also the following section).

Before proceeding further, we point out that although the formulation of the diffusion theory in Sec. 2.1 and Sec. 2.3 is different, as shown in Appendix E, a power series expansion in *λ*(*n*) of (11) exactly matches the result expected from (4) which is obtained by explicitly calculating the relevant correlation functions.

### 3.2 Fixation time

We now consider the mean and variance of the fixation time, conditioned on the eventual fixation of the mutant. By differentiating the expression in (11) for the cumulative distribution of the fixation probability with respect to time, we obtain the probability that the mutant fixes at time *t* using which the *m*th moment of the conditional fixation time can be found. On averaging over the initial population size, we obtain

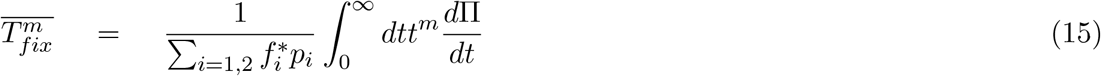

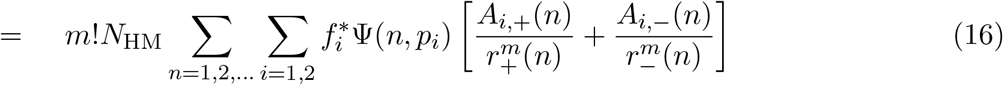

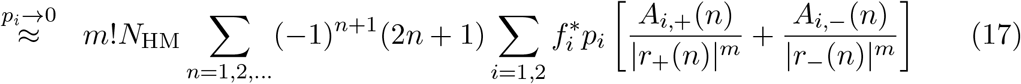

since for small *p*, Ψ(*n, p*) ≈ *p*(−1)^*n*^(2*n* + 1). For *m* = 1, after some algebra, we obtain

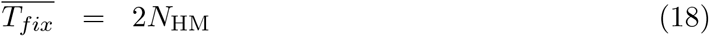

Thus, the conditional mean fixation time of a rare neutral mutant in an expanding and contracting population is simply given by that in a constant population [28] of size *N*_HM_. For given *N*_1_, *N*_2_ and 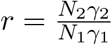, as 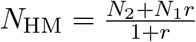, the conditional mean fixation time is independent of *N*_1_*γ*_1_, and thus (18) holds for any rate of demographic fluctuations; this result is also consistent with the observation in the last section that the long time behavior of the average fixation probability depends weakly on *N*_1_*γ*_1_.

However, the conditional mean fixation time, given that the mutant arises in the population of initial size *N*_*i*_, varies with *N*_1_*γ*_1_. Analogous to (16), we can write

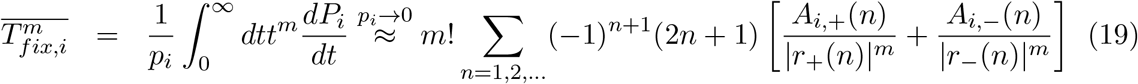

which yields

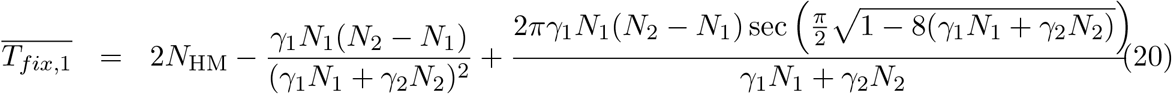

and 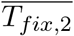 on interchanging the subscripts 1 and 2 in the above expression. Figure 2a shows that 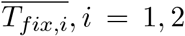 vary monotonically with *N*_1_*γ*_1_, and lie between 2*N*_*i*_ and 2*N*_HM_, as expected [refer to the discussion below (13)] for slowly and rapidly fluctuating population size, respectively. We note that the conditional mean fixation time of the mutant arising in the larger (smaller) population is underestimated (overestimated) by the corresponding result in a model with constant population size *N*_HM_ with the discrepancy increasing with increasing *N*_2_/*N*_1_ (and decreasing *N*_1_*γ*_1_).

**Figure 2:**
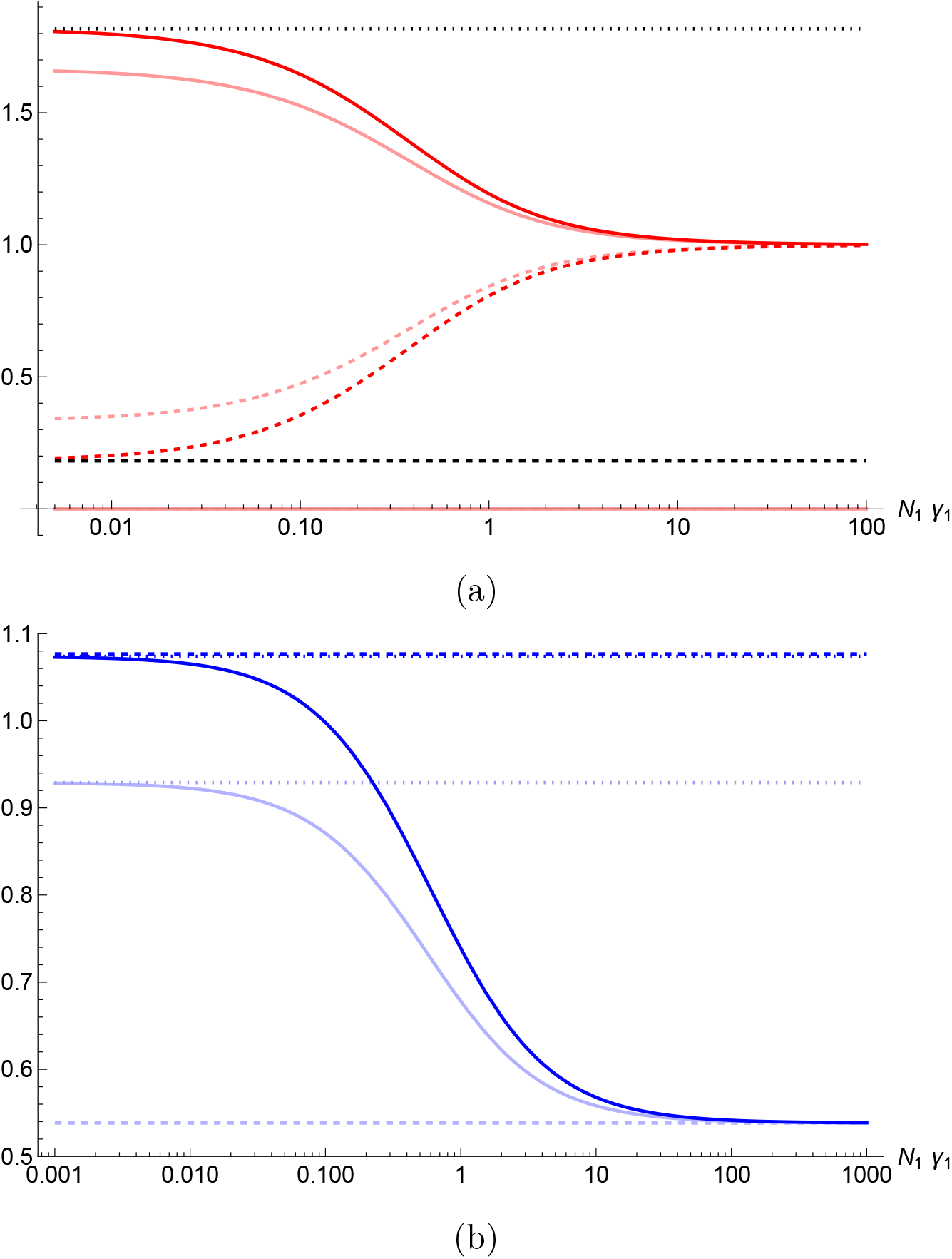
(a) Relative conditional mean fixation time 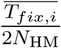, of a mutant that arises in the population of size *N*_1_ (dashed) and *N*_2_ (solid) which are obtained from (20) for *N*_2_ = 5*N*_1_ (light red) and *N*_2_ = 10*N*_1_ (dark red). The relative conditional mean fixation time 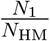 (black dashed) and 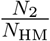 (black dotted) for slowly fluctuating population size are also shown for *N*_1_ = 10^3^, *N*_2_ = 10*N*_1_.(b) Relative standard deviation 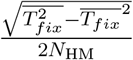 of conditional fixation time averaged over the initial population size which is obtained using (F.2) and (18) for *N*_2_ = 5*N*_1_ (light blue) and *N*_2_ = 10*N*_1_ (dark blue). The dotted and dashed lines, respectively, show the results obtained from (21a) and (21b) for slowly and rapidly fluctuating population size. In all the figures, *N*_1_ = 10^3^ and *N*_2_*γ*_2_ = *N*_1_*γ*_1_.

In a neutral population of constant size, the fluctuations about the conditional mean fixation time increase linearly with the population size [28], and one may ask if this pattern holds in a fluctuating population. From (17), the second moment of the fixation time when averaged over the initial population size can be obtained, and the exact expression is given by (F.2). Using this result and (18), the standard deviation in the conditional fixation time relative to its mean can be found and is shown in Fig. 2b for two values of *N*_HM_. We find that this ratio tends to a constant (in *N*_HM_) only when the environment fluctuates rapidly, and the fluctuations about the mean are larger than expected from a model with constant size *N*_HM_ when the population size fluctuates slowly or moderately fast. Thus, perhaps surprisingly, the fluctuations in the conditional mean fixation time are smaller when the population size fluctuates rapidly. For slowly and rapidly fluctuating population size, using the results in Appendix F, we find that the variance in the conditional fixation time is given by

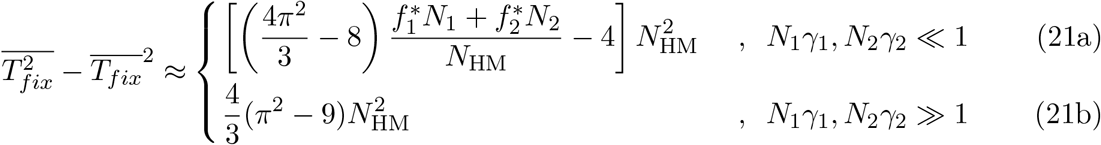

Since the first and second moment of the unconditional fixation time of a single mutant in a constant population of size *N* are given, respectively, by 2 and 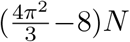 [28], the above results are consistent with the expectation that the variance in the slowly fluctuating environment is a weighted average of the corresponding results in a constant population with size *N*_1_ and *N*_2_, while in the rapidly fluctuating environment, it is given by the result for the population of size *N*_HM_.

### 3.3 Mean sojourn time

In a panmictic, haploid population of constant size *N*, the mean time spent by a neutral mutant between allele frequencies *x* and *x* + *dx* before it gets absorbed decreases as 2/*x* for *N* ^−1^ < *x* < 1 [52, 27]. We now investigate how the fluctuating population size affects the mean sojourn time. For this purpose, we consider (7) and (8) for 0 < *x* < 1 with boundary conditions, *P*_*i*_(0, *t*; *x*) = *P*_*i*_(1, *t*; *x*) = 0 and initial condition, *P*_*i*_(*p*, 0; *x*) = *δ*(*p* − *x*) since the mutant can not reach a frequency 0 < *x* < 1 starting from 0 < *p* < 1 in zero time unless its initial frequency is *x*. For these boundary and initial conditions, a calculation similar to that in Appendix C gives

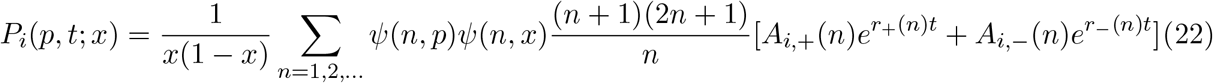

where, as before, *ψ*(*n, p*), *r*_±_(*n*) and *A*_*i*,±_(*n*) are given, respectively, by (C.4), (C.12) and (C.14). From the above expression, the mean sojourn time of a mutant, given that it arises in the population of size *N*_*i*_ is found to be

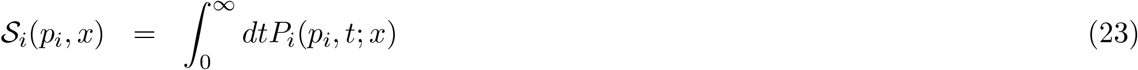

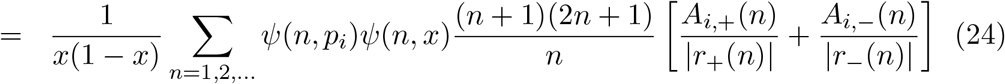

Then the mean sojourn time averaged over initial population size is given by

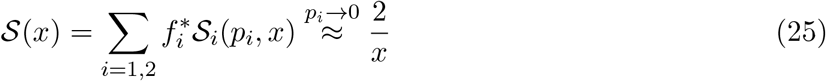

for any rate of demographic fluctuations.

However, using the results in Appendix D for slowly and rapidly fluctuating population size in (24), we find that the mean sojourn time, S_*i*_(*p*_*i*_, *x*), *i* = 1, 2 depend on *N*_*i*_*γ*_*i*_ as

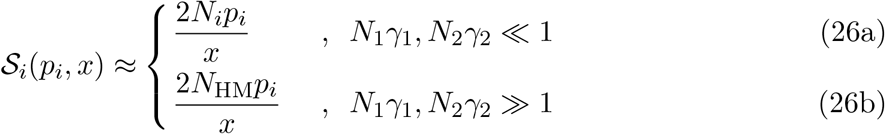

Figure 3a shows the mean sojourn time 𝒮_*i*_(*p*_*i*_, *x*), *i* = 1, 2 for moderately fast fluctuating population size, and we find that 𝒮_*i*_(*p*_*i*_, *x*) lie between the expressions given in (26a) and (26b) but these sojourn times are not proportional to 1/*x*. Using (24) for an initially rare mutant, we obtain

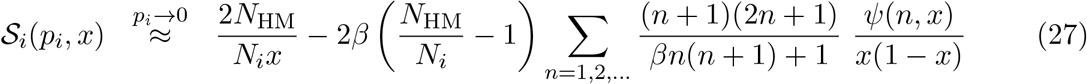

where 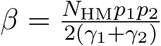.Although the sum in the second term on the RHS of the above equation does not seem to be exactly solvable, as explained in Appendix G, an approximate expression for it can be obtained which finally yields

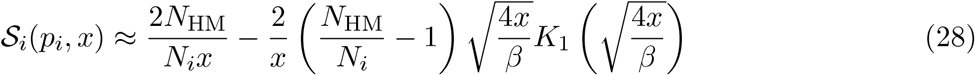

where *K*_*n*_(*x*) is the modified Bessel function of the second kind. In Fig. 3b, the ratio 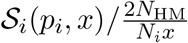 is compared with (28), and we find an excellent agreement.

**Figure 3:**
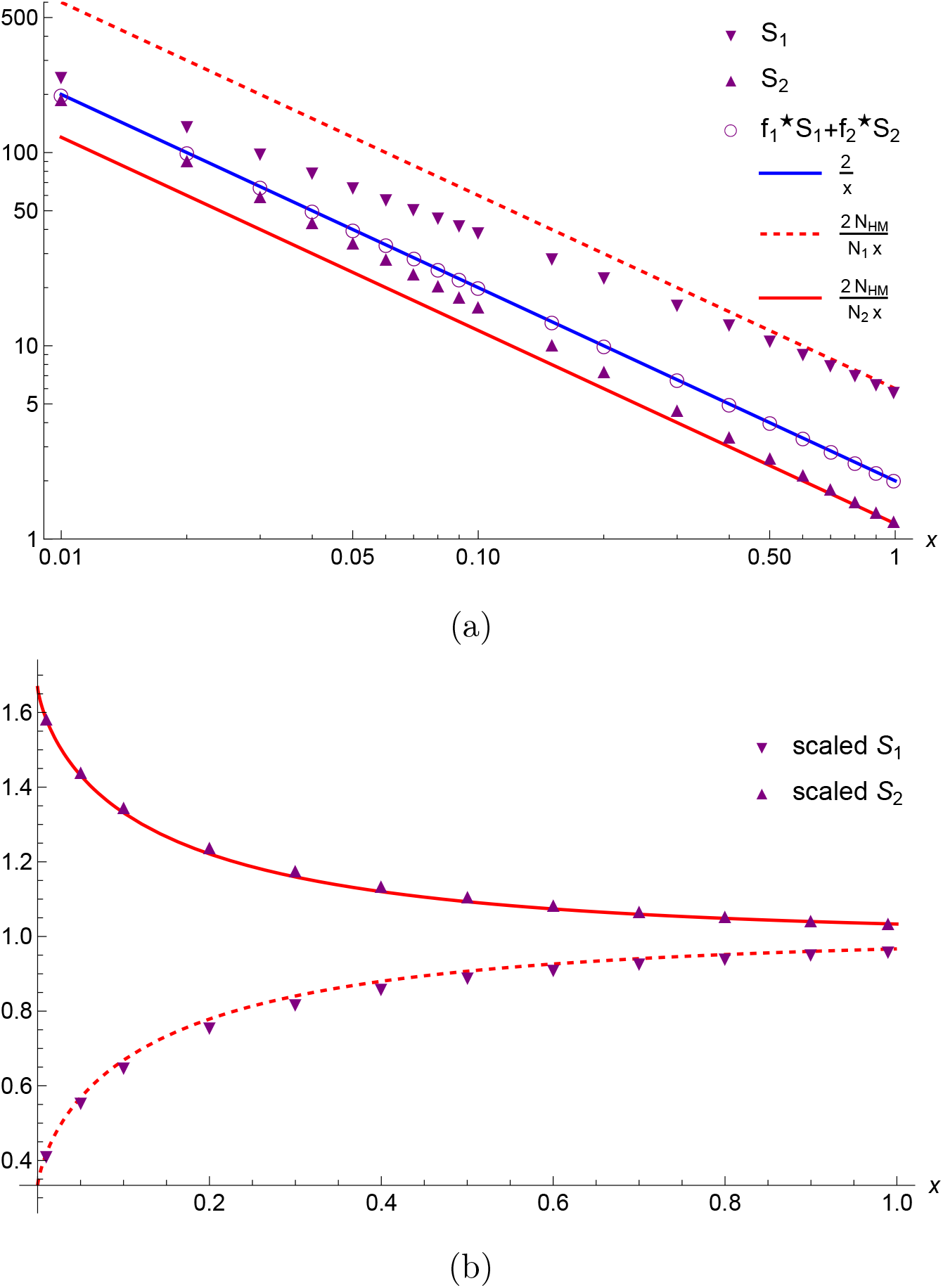
(a) Mean sojourn time 𝒮_*i*_(*p*_*i*_, *x*) of a mutant that arises in a population of size *N*_*i*_, *i* = 1, 2 and (b) scaled mean sojourn time 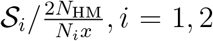 as a function of mutant allele frequency *x*. In both the figures, the parameters are *N*_1_ = 10^3^, *N*_2_ = 5*N*_1_, *N*_2_*γ*_2_ = *N*_1_*γ*_1_ = 1, and the points are obtained numerically from (24). In figure (b), the solid and dashed curves are obtained from (28).

To understand the expression in (28), using (G.3), we obtain

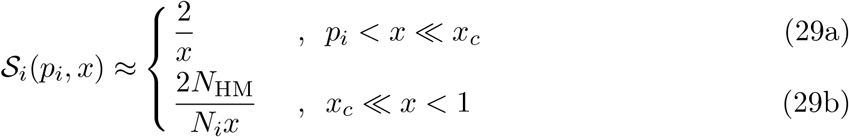

for any rate of demographic fluctuations. In the above expression,

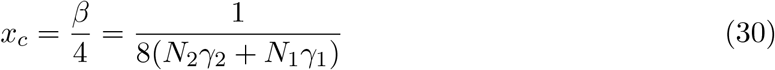

is the allele frequency below and above which the amplitude of the mean sojourn time is different. When the allele frequency is small (*x* ≪ *x*_*c*_), as the mutant has spent a short time in the population, it is not sensitive to the changes in the population size and therefore, we expect the mean sojourn time to be given by that for a single mutant in a population of constant size *N*_*i*_; this expectation is indeed consistent with (29a) as well as (26a) for slowly fluctuating population size. On the other hand, for *x* ≫ *x*_*c*_, due to (30), *Nγ* ≫ *x*^−1^ > 1 and therefore, the large allele frequency behavior for any rate of demographic fluctuations coincides with (26b) for rapidly fluctuating population size.

If one assumes that the effect of fluctuating population size can be subsumed in the effective population size given by *N*_HM_, the mean sojourn time of the mutant with initial frequency *p*_*i*_ will be given by 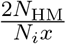 for all *p*_*i*_ < *x* < 1. But, as Fig. 3b shows, if the mutant arises in the larger (smaller) population, the mean sojourn time is underestimated (overestimated) by the constant population size model at low to intermediate allele frequencies although the agreement is good at large allele frequencies.

On integrating both sides of (24) over the mutant allele frequency 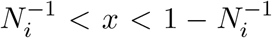,the mean absorption time of a single mutant given that it arises in the population of size *N*_*i*_ can be obtained, and is discussed in Sec. S2 of Supplementary Information. Here, using (26a) and (26b), we note that the mean absorption time is approximately given by 2 ln *N*_*i*_ and 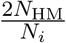 ln *N*_*i*_, respectively, in slowly and rapidly fluctuating environment. This yields the mean absorption time averaged over the initial condition to be 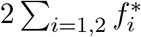 ln *N*_*i*_ for *N*_1_*γ*_1_, *N*_2_*γ*_2_ ≪ 1, and 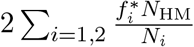 ln *N*_*i*_ for *N*_1_*γ*_1_, *N*_2_*γ*_2_ ≫ 1 which suggests that, unlike the average conditional mean fixation time [see (18)], the mean absorption time is not a constant in *N*_1_*γ*_1_ but, as Fig. S2 shows, the dependence on the rate of demographic fluctuations is rather weak.

## 4 Discussion

Effective population size is an important and useful concept in population genetics as it allows one to subsume some of the details of the population into a single parameter [46]. However, it has also been realized that in a neutral population, when the population size fluctuates on the same time scales as that over which the random genetic drift operates, an effective population size can not be defined [36, 22, 19, 41, 45] and therefore, one can not appeal to the neutral theory for constant population size [26] which raises the question how the classical results are affected in such parameter regimes.

Although numerical studies have been conducted in various specific situations to fill this gap in our understanding [41, 51, 34], an analytical formulation of the problem as described in Sec. 2.1 is useful to draw some general conclusions: first, as discussed below (5), the precise mathematical meaning of the phrase ‘effective population size exists’ is that the exponent in (4) is linear in the eigenvalue *λ*(*n*) (this remark holds when other evolutionary forces such as selection are also present on replacing *λ*(*n*) by the eigenvalue of the corresponding constant environment problem). Second, equation (4) also shows that the biological reason for the nonexistence of *N*_*e*_ lies in the fact that the population sizes are, in general, correlated in time, and it is only in the parameter regimes where these correlations can be ignored, an effective population size can be defined. Third, one can obtain an insight in the parameter range(s) where the effective population size can serve as a good approximation by considering a two-time correlation function. For example, here we showed that the correlation between the inverse population size at two different time points decays exponentially at the rate given by the sum of the switching rates [refer to (B.10) and (E.6)]. Then, if the time is scaled by one of the population sizes, the correlations become negligible when the product of the population size and switching rates is either small or large compared to one.

We also considered a simple demography model in which the population size switches randomly between two values, and obtained exact expressions for various quantities of interest. It is instructive to compare these results with a neutral model (henceforth referred to as *N*_*e*_ model) which assumes that the effect of fluctuating population size can be absorbed in a constant effective population size given by the harmonic mean of the population sizes; as discussed below (5), this assumption corresponds to setting correlations between the population size at different times to zero. We find that the predictions from these two models differ significantly when the switching times are similar to the population size in the full model; for example, as shown in Fig. 1, the fixation probability of a mutant at short times is underestimated by the *N*_*e*_ model.

In a recombining population, the conditional mean fixation time plays an important role in determining genetic diversity when a selective [3] or neutral [33] sweep occurs, and it has been shown that longer the fixation time (relative to recombination time), smaller the genetic diversity at the end of the sweep. Our results show that when the population size fluctuates at moderately fast rate, the fixation time of a mutant that arises in the larger population is larger than that predicted by the *N*_*e*_ model [refer to Fig. 2] which suggests that the latter model overestimates the neutral genetic diversity in these parameter regimes. A similar conclusion is drawn for the mean sojourn time which, at intermediate mutant frequencies, is not proportional to the reciprocal of the allele frequency [52, 27], and is underestimated by the *N*_*e*_ model for a mutant arising in the larger population [see Fig. 3].

These results suggest that in general demographies, the neutral genetic diversity is likely to be quite different from the predictions from the *N*_*e*_ model when the relevant time scales are not well separated, and further studies, perhaps in the framework of the infinite-sites model, are desirable for a deeper understanding of the effect of demography on the measures of neutral genetic diversity [4]. An important caveat of our work is the assumption of neutrality - although we expect that these results will continue to hold, at least qualitatively, for weak selection, it is an open question to understand how genetic drift, demographic fluctuations and strong selection interact to affect the dynamics of an evolving population.

## Appendix A Discrete time simulations

In the simulations, the total population size and the number of mutant allele evolve in discrete time. We assume that the population size takes two values *N*_1_ < *N*_2_: if the population size in the current generation is *N*_1_, it either changes to *N*_2_ with probability 0 < *q*_1_ < 1 or remains unchanged with probability 1 − *q*_1_ in the next generation; similarly, the population size *N*_2_ → *N*_1_ with probability *q*_2_ or remains *N*_2_ with probability 1 − *q*_2_. At long times, when this two-state Markov chain reaches a stationary state, the population dynamics are implemented using a Wright-Fisher process as described in the main text (see Sec. 2.2).

The model described above was also compared with the one in which when the population size switches, the number of mutants in the next generation are not stochastically sampled but are given by 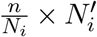 (with *N*_*i*_ and 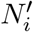,respectively, being the population sizes at current and next generation), which keeps the mutant *frequency* unchanged immediately after the switch, otherwise the standard Wright-Fisher process for constant population size is implemented. Both the models were simulated starting with a single mutant, irrespective of the population size in which the mutant arises (that is, the initial frequency of the mutant is different). Our numerical simulations showed the results obtained from these updating protocols to be in close agreement.

## Appendix B Random telegraph process and its properties

In the random telegraph process, the probability *f*_1_ and *f*_2_ that the population size at time *t* is, respectively, *N*_1_ and *N*_2_ evolves according to

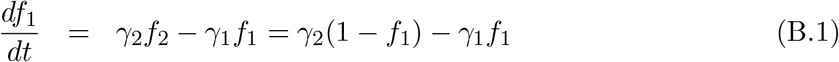

where *γ*_1_ is the rate at which *N*_1_ → *N*_2_ and *γ*_2_ is the reverse rate, and *f*_1_ + *f*_2_ = 1.

In the stationary state where the LHS of the above equation is zero, the probability of the population size being *N*_1_ is

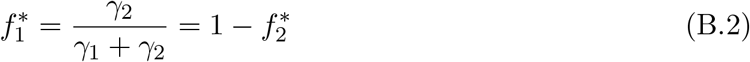

The *n*th cumulant *κ*_*n*_ of the inverse population size in the stationary state can be obtained using the cumulant generating function [29],

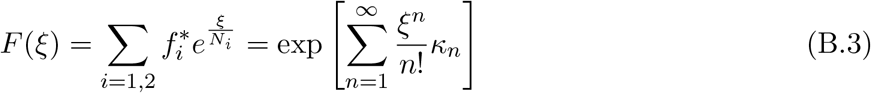

The first few cumulants are then given by

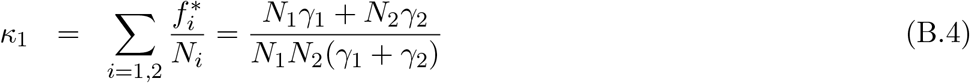

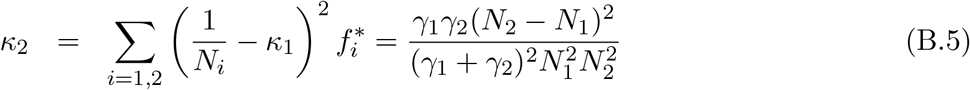

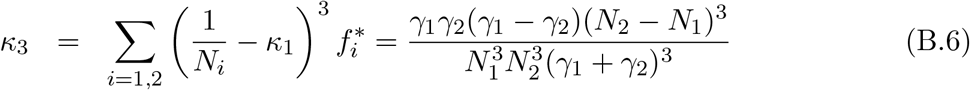

Note that the variance *κ*_2_ and skewness *κ*_3_ vanish when *N*_2_ = *N*_1_, as expected, and their magnitude increases with the difference in the population sizes.

Equation (B.1) for the dynamics of *f*_*i*_(*t*) can be easily solved to give

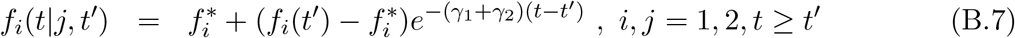

where *f*_*i*_(*t*^′^) = *δ*_*j*,*i*_ is one when given the population size at *t*^′^ is *N*_*i*_ otherwise it is zero. Using the above equation and on averaging over the initial population size (chosen from stationary telegraph process), we obtain the mean of the inverse population size to be

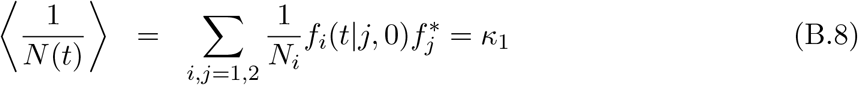

which is independent of time, as expected in the stationary state. Similarly, the (two-point) correlation function is given by

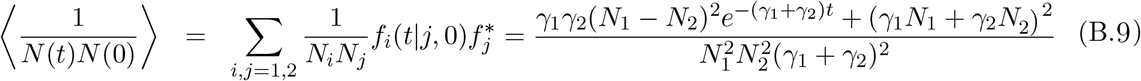

which yields

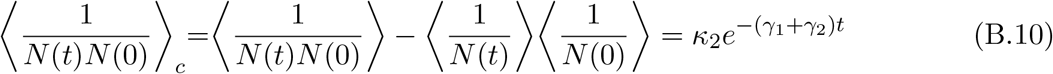

## Appendix C Exact solution of the fixation probability

Here, we obtain an exact solution of (7) and (8) subject to boundary conditions (9). As the boundary condition at *p* = 1 is inhomogeneous, we write *P*_*i*_(*p, t*) = *p* + *F*_*i*_(*p, t*), *i* = 1, 2 where, due to (7) and (8), we have

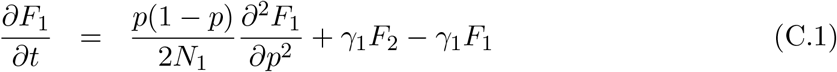

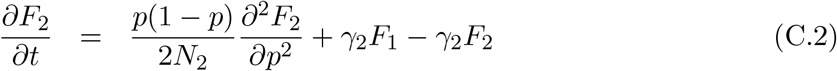

with *F*_*i*_(0, *t*) = *F*_*i*_(1, *t*) = 0. We then expand *F*_1_, *F*_2_ as a linear combination of Jacobi polynomials with time-dependent coefficients,

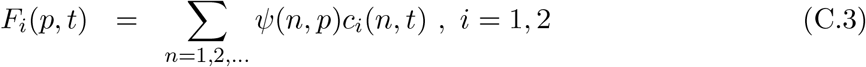

where

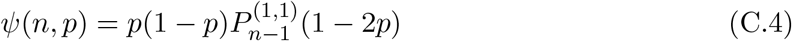

since the Jacobi polynomials obey 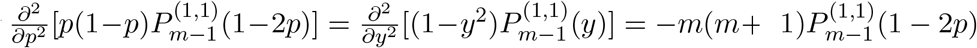. For the initial condition, *P*_*i*_(*p*, 0) = 0 or *F*_*i*_(*p*, 0) = −*p*, using (C.3) at *t* = 0, we obtain

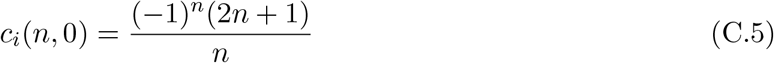

where we have used [see table 18.3.1 of [38]]

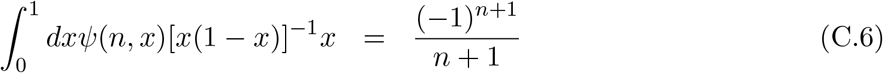

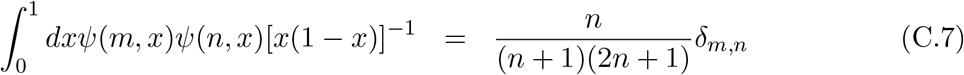

Using the expansion (C.3) in (C.1) and (C.2) and the fact that the Jacobi polynomials are orthogonal, we obtain linear, first order, coupled differential equations for *c*_*i*_(*n, t*):

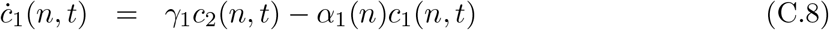

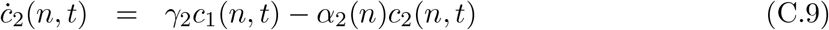

where

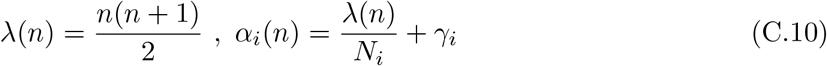

From (C.8) and (C.9), we find that both *c*_1_, *c*_2_ obey 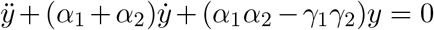 which can be easily solved, and we obtain

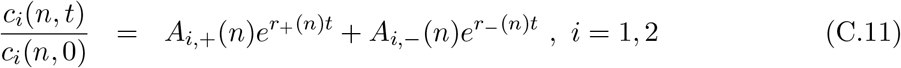

where, *r*_±_ obey *r*^2^ + (*α*_1_ + *α*_2_)*r* + (*α*_1_*α*_2_ − *γ*_2_*γ*_1_) = 0, and given by

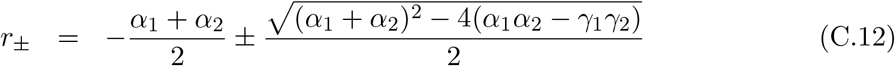

Note that *r*_−_(*n*) < *r*_+_(*n*) < 0. To find *A*_*i*,±_, we first note that *A*_*i*,+_ + *A*_*i*,−_ = 1 to satisfy (C.11) at *t* = 0. Furthermore, the coefficients *A*_1,±_ and *A*_2,±_ are related due to (C.8) or (C.9) as

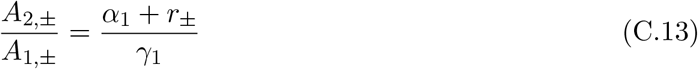

Solving these equations, we finally obtain

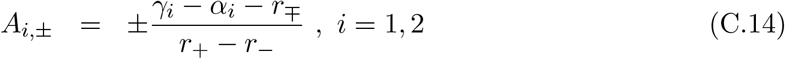

Thus we can write

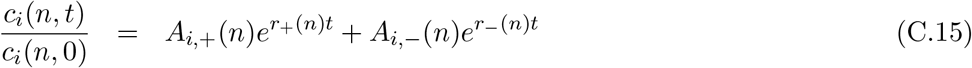

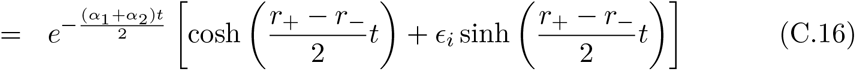

where,

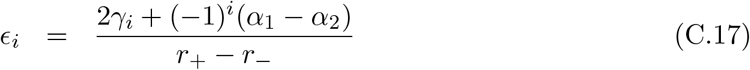

## Appendix D Fixation probability in slowly and rapidly changing environment

Here we analyze the exact expression (11) when 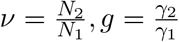 are kept fixed, and *ℓ* = *N*_1_*γ*_1_ are very small or large compared to one. For *ℓ* ≪ 1, on expanding *r*_±_, *A*_*i*,±_ given in Appendix C, about *ℓ* = 0, we obtain

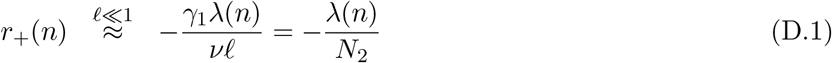

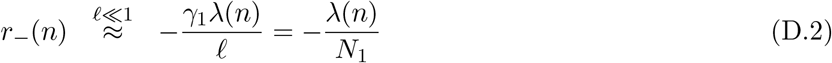

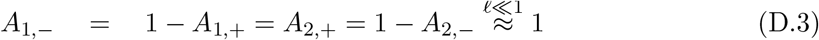

On using these results in (C.3) and (C.11), we obtain

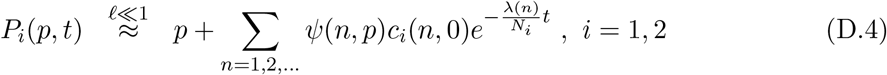

Similarly, when *ℓ* ≫ 1, we have

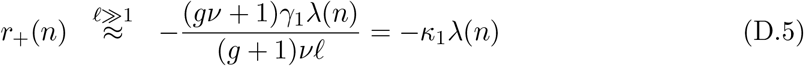

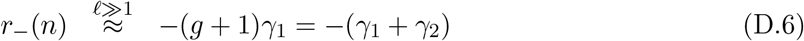

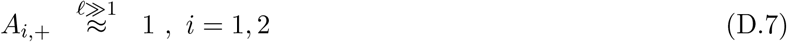

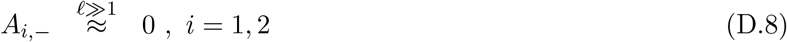

where we have used (B.4). We thus obtain

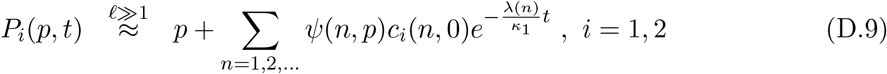

## Appendix E Cumulant expansion of fixation probability

Here we show that a power series expansion of the exact result (11) about *λ*(*n*) = 0 matches the results expected from (4); we demonstrate this for the fixation probability *P*_1_ and carry out this exercise to quadratic order in the eigenvalue *λ*(*n*). From (C.16), we have

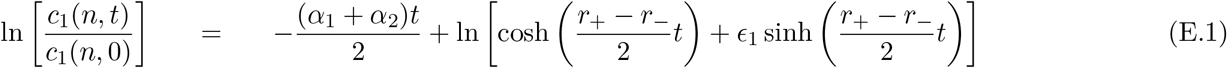

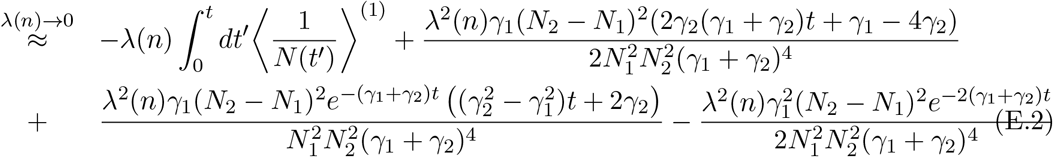

where, 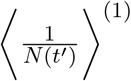 is given below.

On the other hand, if the mutant arises in a population of size *N*_1_, using (B.7), we obtain

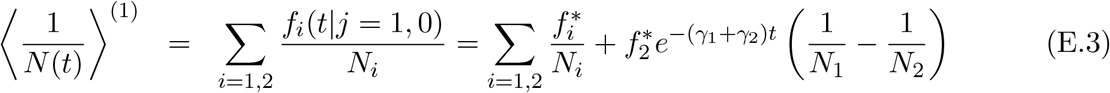

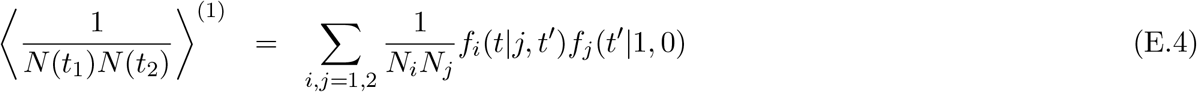

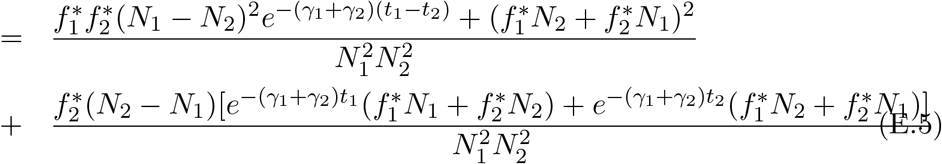

for *t*_1_*>* > *t*_2_ > 0 so that

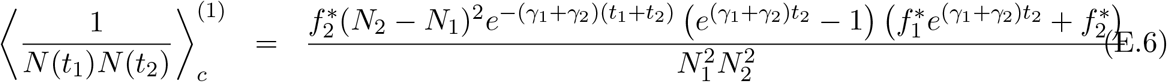

Using these expressions, the integrals appearing on the RHS of (4) can be easily calculated and found to match exactly with (E.2) above.

## Appendix F Moments of fixation time

From (17), for *m* = 2, we obtain

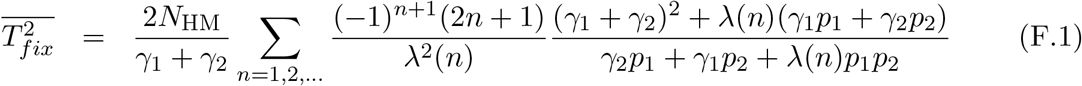

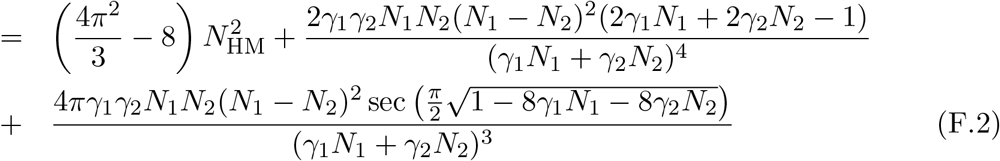

As desired, the last two terms on the RHS vanish when *N*_2_ = *N*_1_ and we are left with the first term which is the result for a constant population [28]. Keeping the ratio of the population sizes and switching rates fixed, the above expression yields

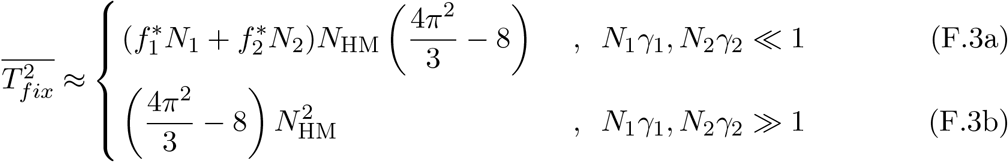

## Appendix G Mean sojourn time

To find an approximate expression for the mean sojourn time 𝒮_*i*_(*p*_*i*_, *x*), we first note that for *m* ≫ 1, *x* ≪ 1 such that *m*^2^*x* is finite, we can write [25],

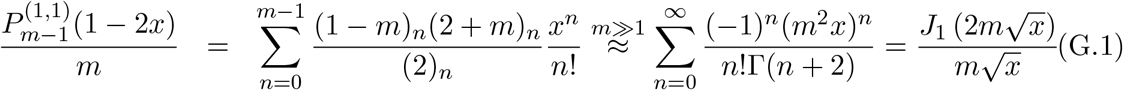

where (*b*)_*n*_ = *b*(*b* + 1)…(*b* + *n* − 1) and *J*_*n*_(*x*) is the Bessel function of the first kind. Then, the sum on the RHS of (27) can be approximated as

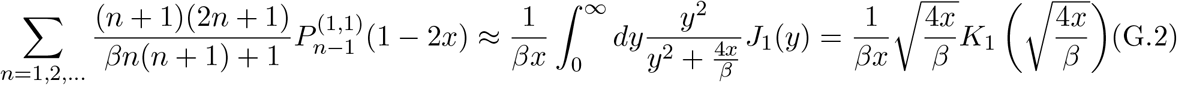

where *K*_*n*_(*x*) is the modified Bessel function of the second kind. Furthermore, since 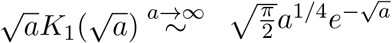 and 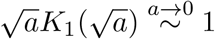,we have

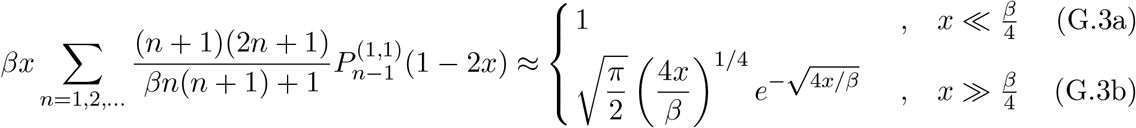

## Supplementary Information

### S1 Fixation probability

**Figure S1:**
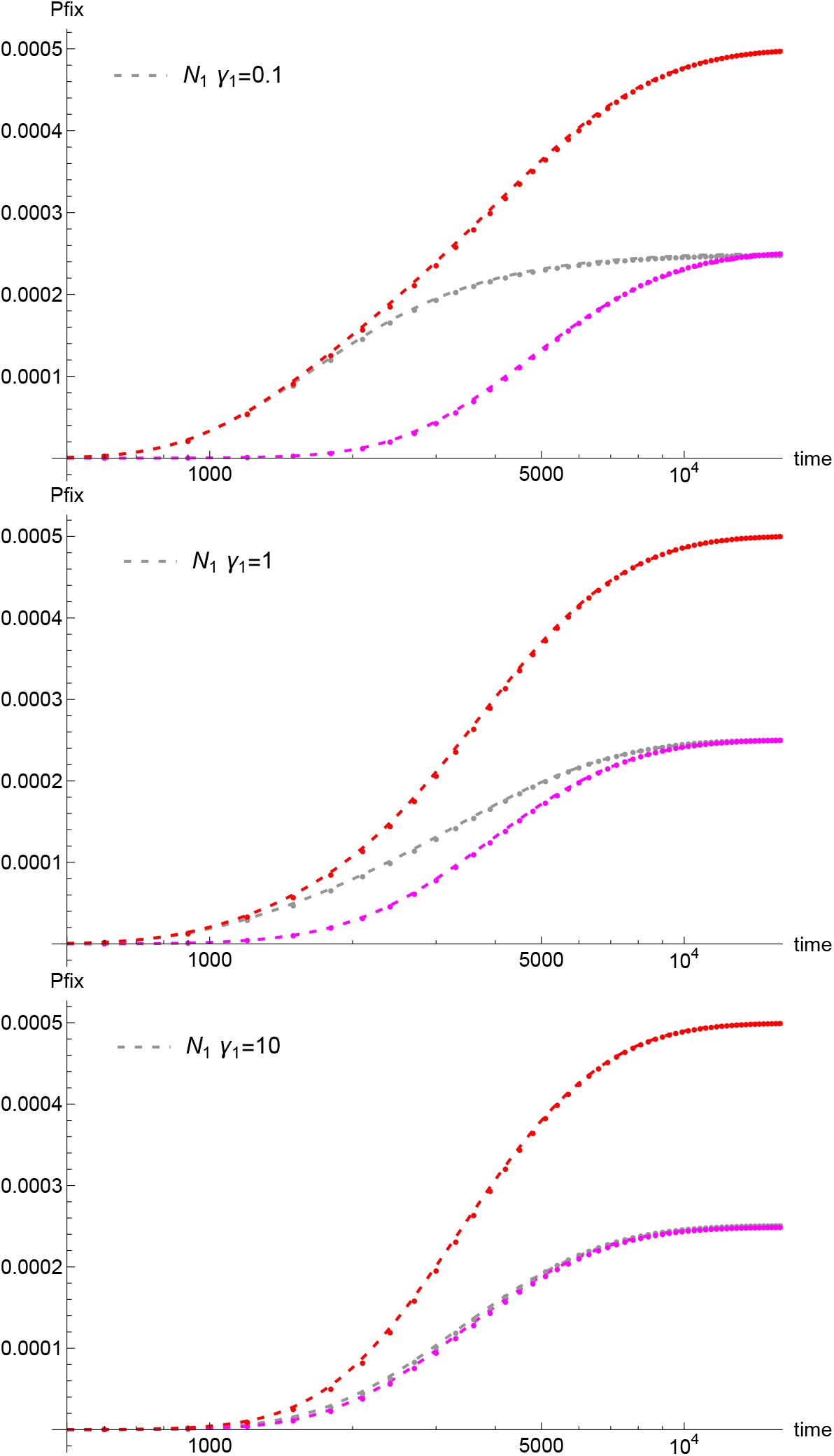
Fixation probability of a single mutant given that it arises in the population of size *N*_1_ (gray) or *N*_2_ (magenta), and when averaged over the initial conditions (red). The simulation results are shown by dots and the numerical solutions of (7) and (8) are given in dashed lines for *N*_1_ = 1000, *N*_2_ = 3000 with *N*_1_*γ*_1_ = *N*_2_*γ*_2_ = 0.1, 1, 10. The simulation data are obtained by averaging over 5 × 10^4^ independent trajectories of the population size.

### S2 Mean absorption time

The probability 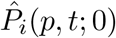 that a mutant arising in a population of size *N*_*i*_, *i* = 1, 2 is lost by time *t* also obeys (7) and (8) in the main text; on solving these equations with boundary conditions, 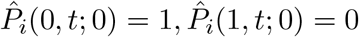 and initial condition, 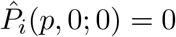,we obtain

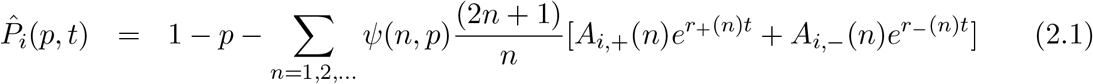

which gives the unconditional mean extinction time to be

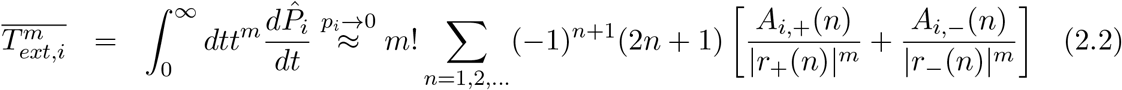

Then the mean absorption time of the mutant arising in frequency *p*_*i*_ in population of size *N*_*i*_ is given by adding the unconditional mean fixation and extinction times to give

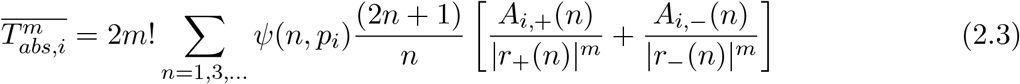

The mean absorption time in a population of constant size *N* when the initial frequency of the mutant is *p* is then given by 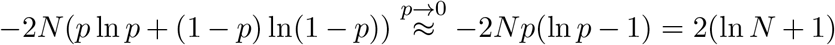 for *Np* = 1 [refer to (5.19) of [10]]. This result can be used to understand the absorption time in slowly and rapidly fluctuating population size as shown in Fig. S2, and discussed in Sec. 3.3 in the main text.

**Figure S2:**
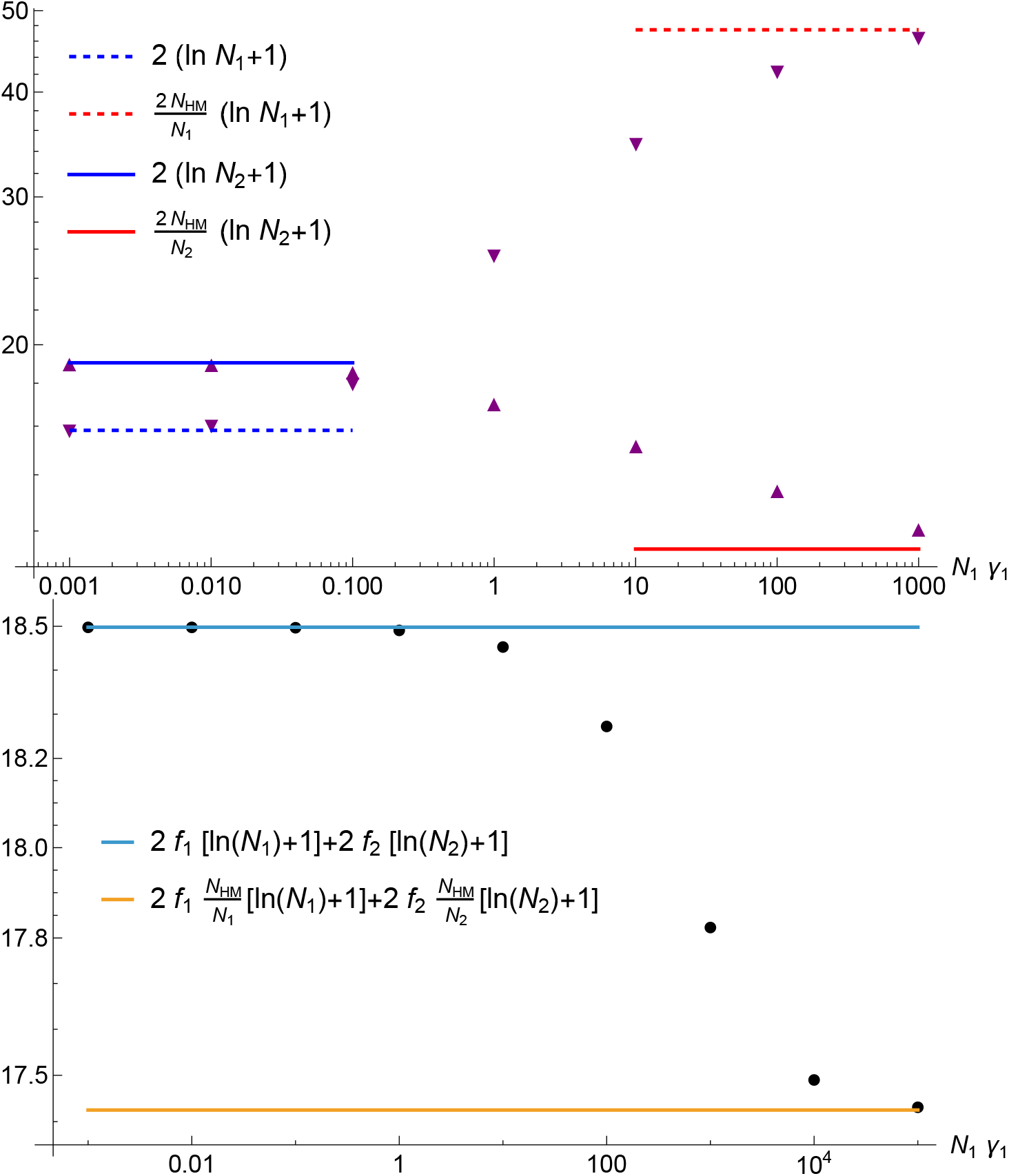
Mean absorption times, 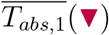 and 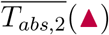 (top) and mean absorption time when averaged over the initial population size, 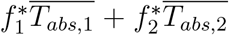 (bottom) as a function of *N*_1_*γ*_1_. The parameters in both the figures are *N*_1_ = 10^3^, *N*_2_ = 5*N*_1_, *N*_2_*γ*_2_ = *N*_1_*γ*_1_, and the points show the results obtained using the exact equation (S2.3).

## Notes

### Competing Interest Statement

The authors have declared no competing interest.

